# Testing Darwin’s hypothesis about the most wonderful plant in the world: The Venus flytrap’s marginal spikes are a ‘horrid prison’ for moderate-sized insect prey

**DOI:** 10.1101/318790

**Authors:** Alexander L. Davis, Matthew H. Babb, Brandon T. Lee, Christopher H. Martin

## Abstract

Botanical carnivory is a novel feeding strategy associated with numerous physiological and morphological adaptations. However, the benefits of these novel carnivorous traits are rarely tested. Here, we used field observations and lab experiments to test the prey capture function of the marginal spikes on snap traps of the Venus flytrap (*Dionaea muscipula*). Our field and laboratory results suggested surprisingly inefficient capture success: fewer than 1 in 4 prey encounters led to prey capture. Removing the marginal spikes decreased the rate of prey capture success for moderate-sized cricket prey by 90%, but this effect disappeared for larger prey. The nonlinear benefit of spikes suggests that they provide a better cage for capturing more abundant insects of moderate and small sizes, but may also provide a foothold for rare large prey to escape. Our observations support Darwin’s hypothesis that the marginal spikes form a ‘horrid prison’ that increases prey capture success for moderate-sized prey, but the decreasing benefit for larger prey is unexpected and previously undocumented. Thus, we find surprising complexity in the adaptive landscape for one of the most wonderful evolutionary innovations among all plants. These findings further enrich our understanding of the evolution and diversification of novel trap morphology in carnivorous plants.

## Introduction

The origins of novel structures remain an important and poorly understood problem in evolutionary biology (Mayr 1960, Mozcek 2008). Novel traits are often key innovations providing new ecological opportunities (Maia et al. 2013; Stroud and Losos 2016; Wainwright et al. 2012). Despite the importance of these traits, our understanding of the adaptive value of novel structures is often assumed, and rarely tested directly. Frequently, this is because it is difficult or impossible to manipulate the trait without impairing organismal function in an unintended way; however, many carnivorous plant traits do not present this obstacle.

Botanical carnivory is a novel feeding strategy that has evolved at least nine separate times in over 700 species of angiosperms, typically in areas with severely limited nitrogen and phosphorus (Ellison 2006; Givnish 2015: Givnish et al. 1984; Król et al. 2012, Roberts and Oosting 1958). Pitfall traps evolved independently at least 6 times and sticky traps 5 times. However, snap traps have most likely evolved only once in the ancestral lineage leading to the aquatic waterwheel (*Aldrovandra vesiculosa*) and Venus flytrap (*Dionaea muscipula*), which is sister to the sundews (Drosera spp.) and within the Caryophalles (Cameron 2002, Givnish 2015, Walker et al. 2017). Multiple hypotheses have been proposed for why snap traps evolved including the ability to capture larger prey, capture prey more quickly, or more completely digest prey (Darwin 1875; Gibson and Waller 2009). However, these hypotheses have never been tested except for a few field studies documenting the size and diversity of arthropod prey (Gibson 1991; Hutchens and Luken 2015; Youngsteadt et al. 2018).

The marginal spikes found in *Dionaea* are modified trichomes that extend from the margin of the trap lobes. These spikes are homologous to the trichomes of sundews, but do not exude any sticky resin and have lost the mucus glands (Gibson and Waller 2009). Darwin was the first to document evidence for carnivory in flytraps and sundews in a series of careful experiments and proposed that the marginal spikes of flytraps enhance prey capture success by providing a cage-like structure around the top of the trap that contains the prey (Darwin 1875; Gibson and Waller 2009). Darwin (1875) also hypothesized that while small insects will be able to escape between the spikes, a moderately sized insect will be “pushed back again into its horrid prison with closing walls” (page 312), and large, strong insects will be able to free themselves. Determining the function of the marginal spikes is important for understanding the rarity of mechanical snap traps whereas sticky and pitfall traps are ubiquitous across carnivorous plants.

Traits that enhance prey capture ability are expected to be strongly selected for given the benefits of additional nutrients and the energetic and opportunity costs associated with a triggered trap missing its intended prey. Nutrients from insect prey increase the growth rate of Venus flytraps (Darwin 1878; Roberts and Oosting 1958) at a cost of lower photosynthetic efficiency of carnivorous plants compared to other plants (Ellison and Gotelli 2009; Pavlovic et al. 2009). The traps are triggered by an action potential when specialized trigger hairs are stimulated (Volkov et al. 2008, 2009) and close as quickly as 100 milliseconds forming a cage around the prey item (Poppinga et al. 2013). If the trap fails to capture an insect, it takes between two and three days for the trap to re-open, during which time it is unable to be used for prey capture. Beyond the energy expended to close a trap and the opportunity cost of a miss, there is a cost associated with declining trap performance and trap death. Traps that have closed and re-opened have lower subsequent trap closure speeds and trap gape angle (Stuhlman 1948). Additionally, after a few closings, traps rapidly die. The marginal spikes provide a novel and unique function that potentially increases prey capture rate and minimizes the costs associated with a failed trap closing event.

We measured prey capture efficiency and the effect of marginal spikes using field observations of wild Venus flytraps and laboratory experiments. By testing the prey capture ability of plants with intact spikes and ones with the spikes clipped off, we assessed the novel function of the marginal spike cage for prey capture.

## Methods

### Field Data Collection

The Green Swamp Preserve, NC, USA is one of the last remaining eastern pine savanna habitats containing endemic flytraps. To estimate prey capture rates, we identified individual plants (n = 14) and recorded the number of traps that fell into four categories: alive and closed, dead and closed, alive and open, and dead and open. All closed traps (*n* = 100) had their length, defined here as the widest point of the lobes on the long axis, recorded with digital calipers. We used a flashlight to illuminate the trap from behind making anything inside the trap visible as a silhouette. If the trap contained something it was assigned a value of 1 for “catch” and if it contained nothing it was assigned a 0 for “miss”. We also noted when a trap was closed on another trap or contained debris inside such as sticks or grass (these were considered a miss; n = 7). Logistic regression in R Studio (R Statistical Programming Group 2018; RStudio Team 2015) was used to determine if trap length had a significant effect on prey capture rate in the field.

### Laboratory prey capture experiments

Plants used in lab experiments were tissue-cultured and purchased from commercial suppliers (bugbitingplants.com; stores.ebay.com/joelscarnivorousplants/). The plants were maintained in 40 liter terraria under high-output fluorescent lighting (14-hour daylight cycle) with 8 cm pots submerged in 1-4 cm of reverse osmosis water at all times. Throughout the duration of the experiments, the plants were kept at ambient temperatures under the lights, ranging from 35° C during the day to 22 C at night), and 50 – 90% humidity. Crickets were purchased from Petsmart and kept in 4-liter plastic containers with shelter, water, and a complete diet (Fluker’s cricket food).

To assess the adaptive role of marginal spikes, we set up prey capture arenas (Fig 1C). Each arena consisted of one plant in a petri dish of distilled water, one cricket of known length (range: 0.7 cm – 2.3 cm) and mass (range: 0.026 g – 0.420 g), cricket food, and a ramp from the dry bottom of the arena to the plant. Only healthy crickets with all six legs were used for prey capture trials. Crickets were chosen as the prey item because they represent one extreme of prey difficulty (large and able to jump) while still making up approximately 10% of the flytrap’s diet in the wild (Ellison and Gotelli 2009). All closed traps were initially marked. We checked the plants for closed traps after three days and after one week. Every closed, empty trap was recorded as a 0 for “miss” and every closed trap that contained prey was recorded as a 1 for “catch”. Following one unmanipulated trial with the spikes intact, we used scissors to clip the spikes from every trap on the plant (Fig 1). The plants were then allowed to recover for a week until the traps re-opened. After the traps re-opened, we placed each plant through a second trial with a new cricket. We performed 51 prey capture trials (34 plants total, 17 used only for unmanipulated trials, and 17 used once before and after spike removal). Only 1 trial resulted in no traps triggered over the full week. We also set up control trials (n = 5) with a newly dead cricket placed on the bottom of the tank and negative controls with no cricket at all (n = 2) to ensure that any experimental trap closures were triggered by the cricket and not spontaneous.

**Figure 1:**
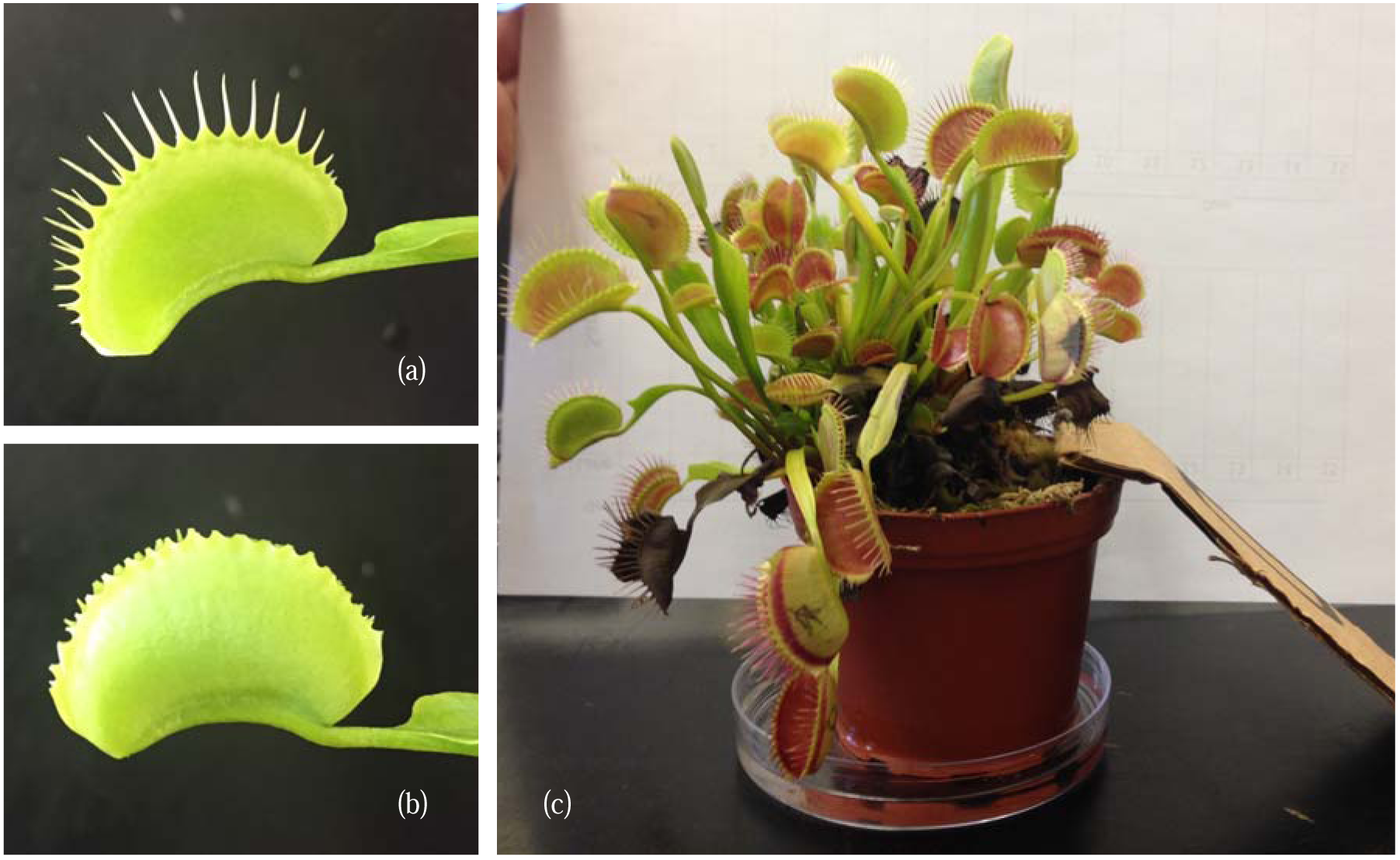
(a) Intact trap; (b) trap with the marginal spikes removed; (c) representative prey capture arena containing one plant, one cricket, a ramp, and a petri dish of water.

To analyze the relationship between prey mass, treatment, trap length, and prey capture success we used multiple logistic regression models in R and generalized linear mixed-effect models using the lme4 package (Bates et al. 2015). For the linear mixed effect models, we used Akaike information criteria with correction for small sample size (AICc) to compare models. We chose prey capture success as our proxy for performance and fitness due to the evidence that the growth rate of flytraps is greatly enhanced by ingesting insect prey (Schulze et al. 2001). We visualized changes in the performance landscape due to removing marginal spikes by estimating thin-plate splines for trials with and without spikes. We fit splines by generalized cross-validation using the Tps function in the Fields package (Nychka et al. 2015) in R (R Core Team (2017).

## Results

### Field Prey Capture Rates

Only 24% of closed wild flytraps contained prey. This number represents a high-end estimate because anything inside the plants was counted as a catch, despite the possibility that the object was a piece of debris instead of an insect or spider. Of the 98 closed traps recorded, 8 were closed around obvious plant debris, and 2 contained identifiable prey (1 ant and 1 spider). 55% ± 5% (mean +/- SE) of wild flytraps were open and alive, therefore able to capture prey.

### Laboratory Prey Capture Rates

Similarly in the lab, only 16.5% of flytraps successfully captured prey out of all closed traps among unmanipulated plants. Only 5.8% of flytraps on these same plants with marginal spikes removed successfully captured prey. Tissue damage due to clipping marginal spikes quickly healed and clipped traps reopened within 4 days; thus, this disparity does not appear to be due to any deleterious effect of tissue damage. Furthermore, no differences in trap closing speeds, health, or growth rates of manipulated traps were apparent. Indeed, marginal teeth began to regrow within approximately one week after removal, suggesting that we underestimated the effect of spike removal on prey capture since spikes were partially regrown by the end of each trial.

Removing marginal spikes reduced the odds of prey capture by 90% relative to unmanipulated traps from the same plant while controlling for prey mass and trap length (effect of manipulation: *P* = 0.002106; linear mixed-effect model relative to model without treatment variable: ΔAIC_c_ = 11). At large prey sizes and large trap lengths this effect disappears (note that spline SE crosses at large prey and trap sizes; Figs. 3b,c).

### Effect of Prey Mass and Trap Length

A linear mixed effect model with prey mass included provided a far better fit to the data than one without (ΔAIC_c_ = 15). In the full model, prey mass was a significant predictor of prey capture success (*P* = 0.000441), with every 0.1 g increase in prey mass corresponding to a 73% decrease in prey capture performance (Fig 3).

Larger trap size also increases the probability of successful prey capture after controlling for prey size, with every 1 cm increase in trap length increasing the odds of prey capture by 2.9-fold (Table 1). Larger trap size increased prey capture success for both manipulated and non-manipulated plants (Fig 3; logistic regression; manipulated: P = 0.02008; non-manipulated: P = 0.003007). A linear mixed effect model including trap length provided a much better fit for the data than one without (ΔAIC_c_ = 31)

**Table 1.** Generalized linear mixed-effect model showing the effect of removing the marginal spikes (manipulation), trap length, and prey mass on prey capture performance (logistic regression). Significant *P*-values are bolded.

| Model Term | Estimate ± SE | *P* |
| --- | --- | --- |
| Manipulation | -2.32 ± 0.75 | **0.002107** |
| Trap Length | 4.74 ± 1.08 | **0.000011** |
| Prey Mass | -13.36 ± 3.80 | **0.000441** |

## Discussion

We provide the first direct test of how prey capture performance is affected by the presence of marginal spikes, trichomes which provide a novel function in Venus flytraps by forming what Darwin described as a “horrid prison”. We found that the marginal spikes are adaptive for prey capture of small and medium sized insects, but not larger insects. In controlled laboratory prey capture trials, 16.5% of trap closures resulted in successful prey capture whereas only 5.8% of trap closures successfully captured prey when marginal spikes were removed (Fig. 2b-c). We found similarly low prey capture rates in the Green Swamp Preserve, one of the natural habitats of the Venus flytrap: fewer than 25% of trap closures resulted in prey capture (Fig. 2a). Furthermore, only about half of the wild traps were open, alive, and available to catch prey. Given the documented tradeoff between photosynthetic efficiency and carnivory and costs associated with maintaining traps (Ellison and Gotelli 2009; Pavlovic et al. 2009), it is possible that the nutrients acquired from a relatively small number of traps are sufficient to maintain the plant. In support of this hypothesis, other carnivorous plants (*Sarracenia purpurea* and *Darlingtonia californica*) sustain themselves with prey capture rates as low as 2% for ants and wasps, respectively (Newell and Nastase 1998; Dixon et al., 2005). Alternatively, prey capture rates for tropical pitcher plants (*Nepenthes rafflesian*) may reach 100% for ants (Bauer et al. 2008). Given that Venus flytraps fall in the middle of this range for pitfall traps, additional factors beyond prey capture rate may underlie the origins of mechanical snap traps.

**Fig. 2:**
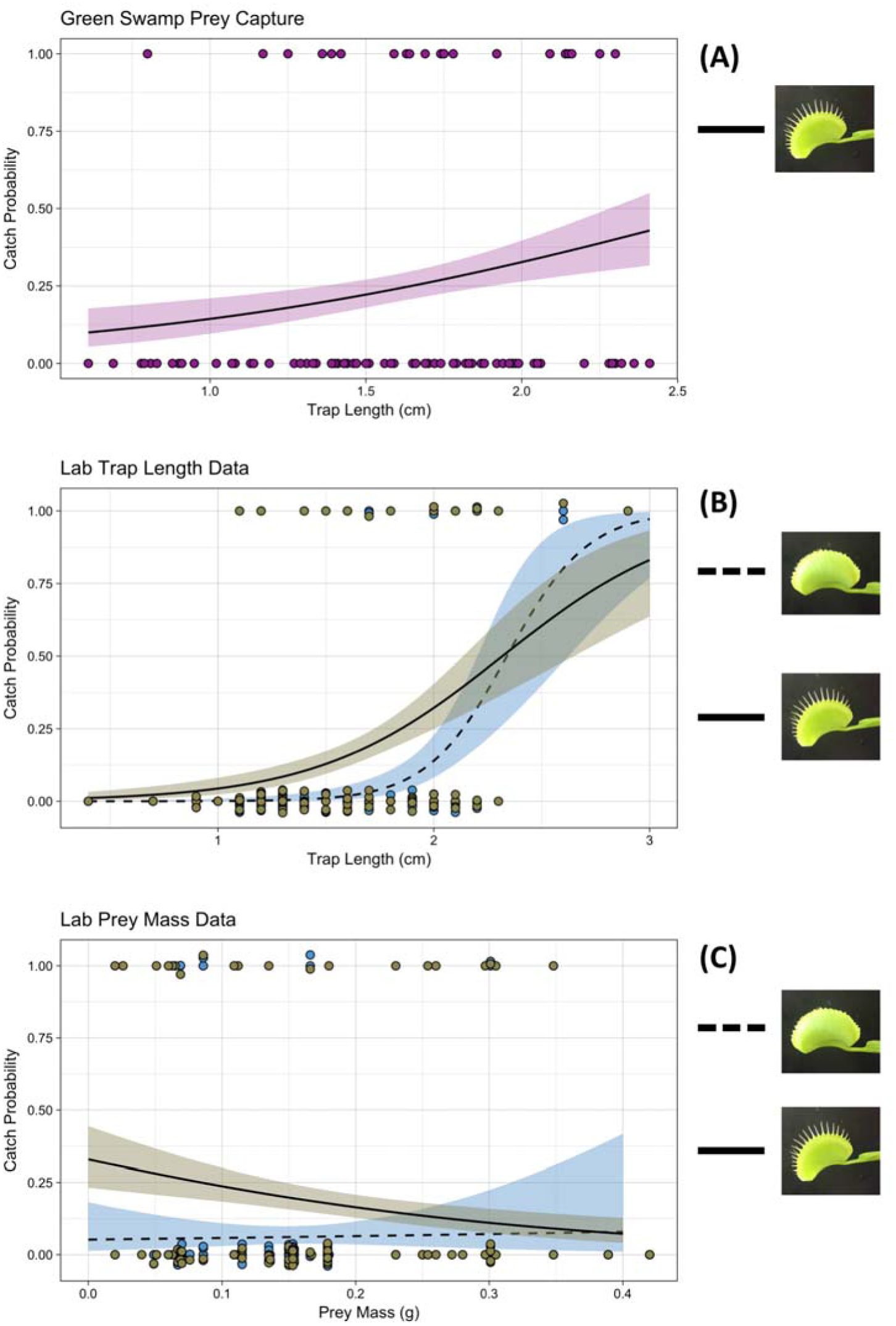
**(A)** Prey capture success of wild plants in the Green Swamp Preserve, NC as a function of trap length (measured to the nearest 0.01”). **(B)** Prey capture success of laboratory plants as a function of trap length (measured to the nearest 0.1”) **(C)** Prey capture success of laboratory plants as a function of prey mass. Lines of best fit were estimated using logistic regression with shaded areas corresponding to ± 1 SE. Each point represents one successful (1) or unsuccessful (0) capture by a flytrap, often resulting in multiple failed captures per cricket mass.

**Fig. 3:**
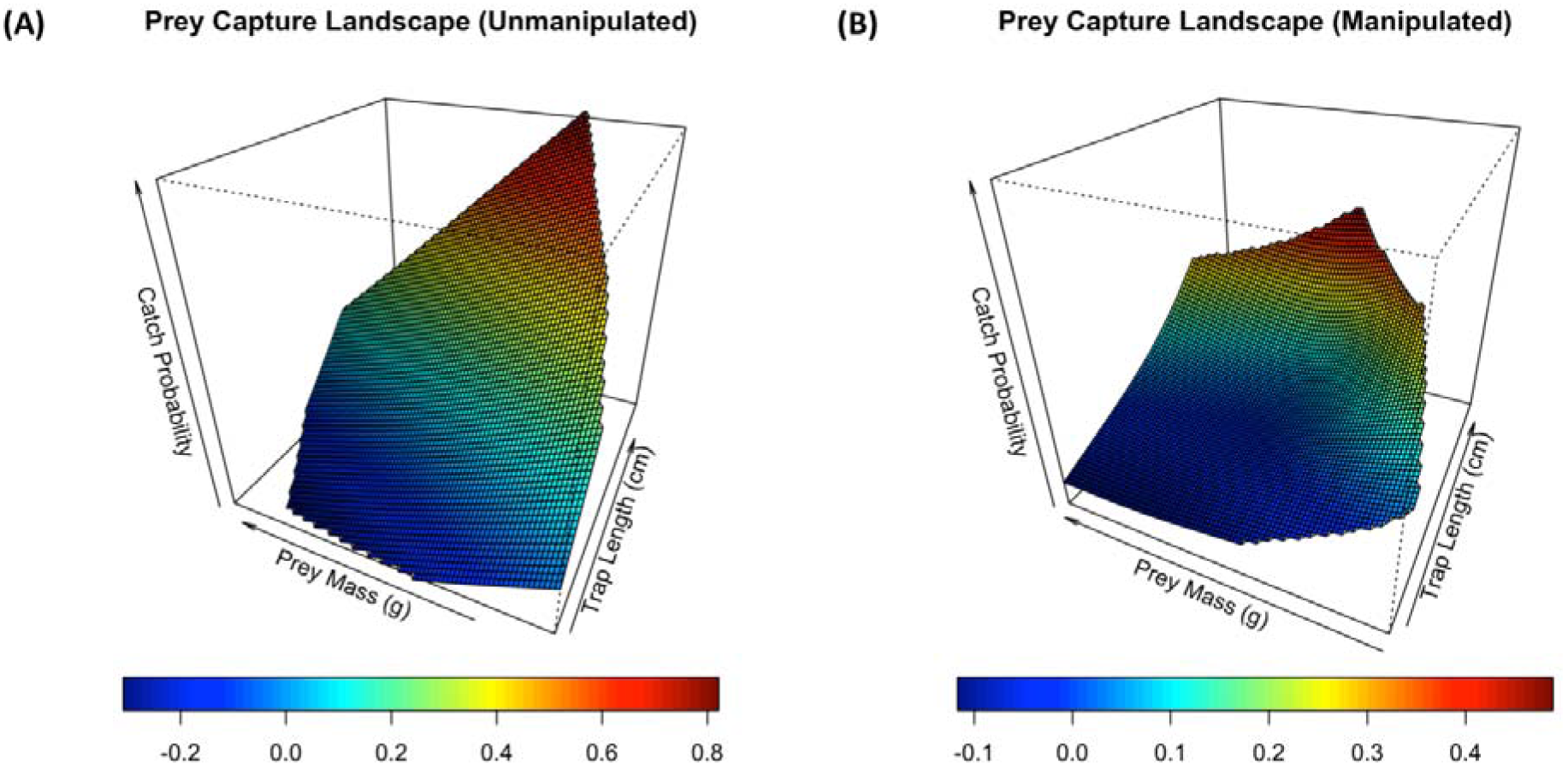
Prey capture landscapes for intact plants (left) and manipulated plants (right). Catch probability is on the z axis and represented by the heat colors relative to insect prey mass and trap length plotted in the x-y plane. The performance landscape for plants without marginal spikes is greatly depressed at small trap sizes, but is similar at large trap/prey sizes.

A second hypothesis for the evolution of mechanical snap traps is selection for capturing larger prey. In habitats where multiple carnivorous plant species coexist we would expect specialization and ecological partitioning (Schoener 1974). Sundews, which grow in sympatry with Venus flytraps in the Green Swamp Preserve, frequently allow prey items larger than 5mm to escape (Gibson 1991) whereas flytraps have been known to capture prey as large as 30mm with an estimated average of 9.3mm (Jones 1923; Ellison and Gotelli 2009). In this study, we found estimated prey capture rates as high as 80% for the largest flytrap sizes despite the average prey size (15.2 mm) being larger than what was reported by Jones (1923). This suggests that mechanical traps are capable of capturing much larger prey than sticky traps. Although some studies found no support for resource partitioning among sympatric species assemblies of carnivorous plants (Ellison and Gotelli 2009; Verbeek and Boasson 1993), others demonstrated differential prey distributions at the individual plant level and among species (Karlsson et al. 1987; Gibson and Waller 2009; Thum 1986). Given the extreme differences in mean and maximum prey sizes between sticky traps and snap traps, it is likely that resource partitioning at least plays a role in the continued coexistence of sundews with flytraps throughout their limited range.

Surprisingly, the effect of removing the marginal spikes for medium-sized traps on prey capture success nearly disappears for larger traps. We observed a possible mechanistic explanation for this counterintuitive result. Crickets are often climbing on the marginal spikes of large traps, and when they trigger them they are able to push against the marginal spikes to pry themselves free. In contrast, when a cricket triggers a large trap with no spikes, it has nothing to use to free itself. Marginal spikes appear to provide leverage for larger insect prey to escape. There is also a possible physical explanation for the diminishing benefit of the marginal spikes at large trap sizes. Stuhlman (1948) speculated that friction between the marginal spikes may slow down trap closure. Because the contact area over which friction matters is proportional to the length squared, we would expect disproportionally large frictional forces as the length of marginal spikes increases on larger traps.

In his writings on insectivorous plants, Darwin (1875) hypothesized that the marginal spikes allowed flytraps to capture larger insects while letting tiny insects go free. Later work has been mixed on whether snap traps are size-selective (Hutchens and Luken 2009; Hatcher and Hart 2014 (ontogenetic changes)) and we did not find any evidence for size-selection here. For medium and small insects, the cage formed by marginal spikes provided a drastic increase in prey capture rates, a finding that is compatible with Darwin’s original hypothesis. At large prey sizes, however, the symmetry between our findings and his hypothesis begin to break down. We found diminishing returns at larger prey sizes, and while Darwin predicted large insects would break free from traps, the mechanism he outlines is different than the one we observe. We did not find that fully trapped insects were breaking free, as he notes in his book. Instead, we found insects that were partially trapped or trapped perpendicular to the trap’s long axis were the ones to break free, potentially with the aid of the marginal spikes.

We demonstrated that the novel marginal spikes, forming a ‘horrid prison’, are an adaptation for prey capture with nonlinear effects at larger prey/trap sizes. Given the diversity of carnivorous plant traps, from the sticky traps of sundews to the rapid suction traps of bladderwort (Brown et al. 2012), we contend that carnivorous plants offer a rich system for investigating the adaptive value of novel traits, particularly within the context of prey capture. Furthermore, this system lends itself to tractable experimental work carried out by undergraduate researchers. This project was carried out during a one-semester course-based undergraduate research experience (CURE) course taught at UNC, entitled ‘The Evolution of Extraordinary Adaptations’. Characterizing the role of these unique features aids our understanding of potential axes of selection that drive the evolution of different trap types and the rarity of mechanical traps. In turn, this tractable laboratory and field systems offers insights into the origins of one of the most wonderful evolutionary innovations among all plants.

## Acknowledgements

We thank Joseph McGirr for logistical assistance and helpful discussion that benefited this work. This work was supported by the Quality Enhancement Program at the University of North Carolina at Chapel Hill, which funded this course-based undergraduate research experience (CURE). Kelly Hogan kindly facilitated and encouraged the development of this course. We also thank Blaire Steinwand and John Bruno for helpful feedback on course design.

## Data accessibility

All data and R scripts used for this study will be deposited in the Dryad Digital Repository.

